# velotest: Statistical assessment of RNA velocity embeddings reveals quality differences for reliable trajectory visualizations

**DOI:** 10.1101/2025.10.26.683064

**Authors:** Sebastian Bischoff, Pavlin G. Poličar, Soham Mukherjee, Jakob H. Macke, Manfred Claassen, Cornelius Schröder

## Abstract

Contemporary single-cell RNA-sequencing studies typically use RNA velocity analyses to investigate cellular dynamics. Commonly, RNA velocities are assessed with two-dimensional projections. Although it is well established that nonlinear embeddings of gene expression data cannot comprehensively preserve the structure of the original high-dimensional data across scales, there is no tool that systematically assesses and quantifies the faithfulness of *velocity* embeddings. This situation potentially leads to misleading velocity visualizations and putative cell state transitions. To this end, we introduce *velotest*, a cell-level statistical test that quantifies the faithfulness of a velocity embedding with respect to its high-dimensional counterpart. Applying velotest across multiple datasets and embedding methods, we find strong differences in the embedding qualities between embedding methods, datasets, and regions in individual datasets. velotest thus provides an essential validation step and enables researchers to select a suitable visualization of high-dimensional velocities, ensuring that hypotheses drawn from velocity projections are grounded in accurately represented dynamics.

## Introduction

Single-cell RNA sequencing (scRNA-seq) is ubiquitously used for the investigation of tissue-level processes at single-cell resolution and has enabled the reconstruction of complex processes in development (e.g., endocrine lineage differentiation in the murine pancreas [1]), differentiation (e.g., murine erythropoiesis [2]), or immune responses (e.g., SARS-CoV-2 infections [3]), yet it only provides a snapshot in time. RNA velocity [4] leverages measurements of both spliced and unspliced mRNA molecules to estimate the time derivative of gene expression, thereby enabling the inference of each cell’s future state and the reconstruction of the temporal order of the process. RNA velocity estimates are high-dimensional velocity vectors, like gene expression profiles, and are typically assessed via visualizations, such as projections into two-dimensional embeddings, generated by t-SNE [5] or UMAP [6]. However, it is well known that two-dimensional, nonlinear embeddings of hundred-dimensional datasets are not able to preserve all relations and properties of the original high-dimensional data [7–9].

This situation begs the question whether two-dimensional velocity visualizations properly represent the corresponding high-dimensional velocity vector [10–12]. Existing velocity visualization approaches [4, 13–16] do not provide a systematic quality assessment, thereby implicitly assuming equally good representations for all data points.

To close this gap, we introduce a statistical test — *velotest* — that, for each cell, evaluates how well its velocity embedding represents the corresponding high-dimensional velocity vector. This test is performed by comparing the embedded velocity against a null distribution of random velocity vectors (Fig. 1a).

**Figure 1.**
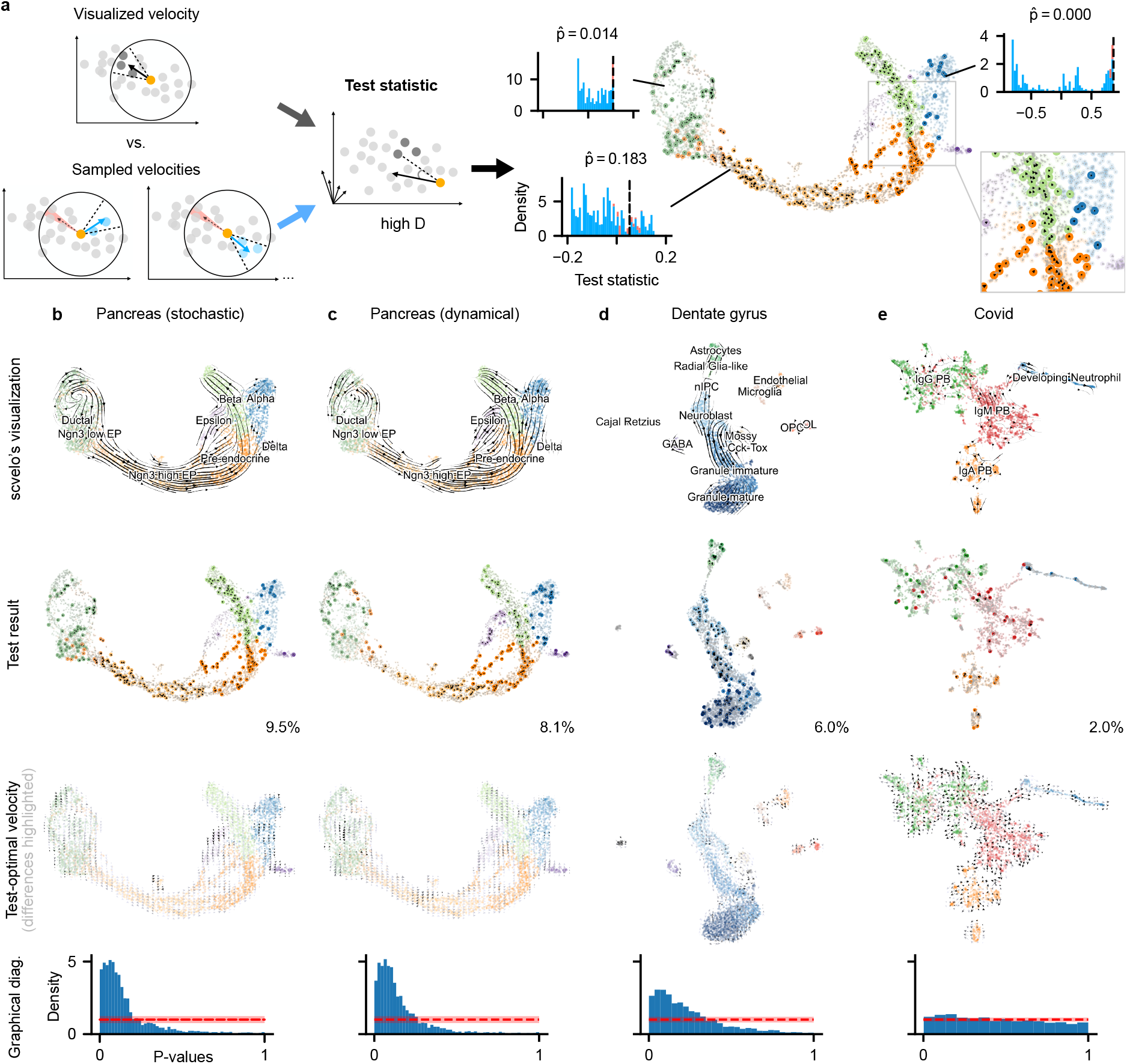
velotest overview and application to scvelo embeddings of multiple scRNA-seq datasets. **(a)** velotest first selects cells in the cone neighborhood (dark grey) of the visualized velocity of the tested cell (orange). Similar neighborhoods are selected for all alternative velocities (light blue), which are outside of an exclusion angle (light red). The test statistics (mean cosine similarity) are computed with the high-dimensional gene expression of the selected cells for the visualized velocities, as well as for alternative velocities. p-values (uncorrected 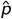 shown) are evaluated according to the observed statistic (black dashed line) and the null distribution originating from the alternative velocities (blue histogram). **(b-e)** scvelo embeddings for pancreas (b-c), dentate gyrus (d), and Covid (e) datasets. Color coding of cells indicates annotated cell types. *First row:* scvelo’s stream plots. *Second row:* visualization of significant tests after correcting for multiple testing (via Bonferroni); cells with significant velocities are colored and velocities shown as black arrows, percentage of significant vectors shown in the lower right corner. *Third row:* test-optimal velocities (grey scale corresponds to differences compared with scvelo’s velocity: deviations over 90°in black, matching velocities in light grey). *Last row:* empirical distribution of p-values with uniform density for the distribution under the null distribution in red (confidence region shaded).

## Results

velotest identifies which cells’ velocities are represented significantly better than chance, highlighting both well- and poorly-represented regions. This allows us to draw conclusions about the overall velocity embedding quality. Furthermore, given a fixed gene expression embedding and our test statistic, we can suggest alternative velocity embeddings, which we call ‘test-optimal velocity’, that more faithfully reflect the high-dimensional velocities. We evaluate velotest for various scRNA-seq datasets, velocity estimation procedures, and visualization approaches. Here, we find only a small subset of velocity visualizations to be significant, as well as situations where velocity visualizations suggest erroneous cellular dynamics. To assess how well the embedded velocities for a scRNA-seq dataset represent their high-dimensional counterparts, we compare the visualized velocity against alternative velocities for each cell and formulate the null hypothesis *H*_0_ that the preimage (i.e., the corresponding high-dimensional velocity vector) of the visualized velocity is no more aligned with the neighboring cells than the preimage of a visually distinct random velocity. Specifically, we compute a p-value for the degree of alignment achieved by the visualized velocity with respect to a null distribution of alignment for alternative velocities (see Methods for details). Multiple testing is accounted for via the Bonferroni correction [17]. Significant velocities are visualized in the original embedding (Fig. 1b-e, second row), indicating which velocities are well-represented in the gene expression embedding, and in turn, which velocities should not be considered for the interpretation of cell state transitions.

We further consider velocity embeddings maximizing the test statistic (test-optimal velocities). These alternative velocities indicate cell-state transitions better aligned with the high-dimensional velocities (Fig. 1b-e, third row). Although these velocity vectors are optimal given the provided gene expression embedding, their interpretation is limited by the quality of the underlying gene embedding. Thus, they may still not accurately represent the high-dimensional velocities (see Fig. S3). We additionally design a graphical diagnostic for an overall assessment across all cells of a dataset. Considering that by definition p-values for non-significant events distribute uniformly, we treat the non-uniformity of the p-value distribution as evidence against the hypothesis that the visualized velocities are random velocity embeddings (Fig. 1b-e, last row).

We apply velotest to multiple scRNA-seq datasets [1–3, 18, 19], and two multi-modal datasets combining scRNA-seq and spatial transcriptomic measurements [20, 21]. Additionally, we assess different embedding methods, namely scvelo [13], veloviz [14], Nyström embeddings [15], and SIRV [22] on the originally analyzed datasets.

We find that scvelo’s [13] visualization of the pancreatic endocrinogenesis dataset [1] accurately represents the overall transition from pre-endocrine cells into beta cells (Fig. 1b, c). However, when we compare embeddings between the stochastic and the dynamic model, i.e. the two most widely used models for estimating the high-dimensional velocity with scvelo, velotest reveals that the visualized transitions from pre-endocrine to epsilon cells for the stochastic model are not supported by the data. In contrast, embedded velocities from the dynamical model support this transition, as indicated by numerous significant velocity vectors (Fig. 1c), and high agreement with the test-optimal velocities (Fig. 1c, Fig. S3a,b). Additionally, we observe that the velocities for the embedding of the stochastic model appear to merge from the epsilon into the beta cluster. This biologically unexpected transition, is not detected as significant by velotest and the test-optimal velocities support the inverse direction. Therefore, we conclude that the unexpected transition stems from a misleading embedding and not from artifactual high-dimensional velocities. Interestingly, the velocity embeddings for the dynamic model support the expected transition and are consistent with the test-optimal velocities. This finding highlights the value of test-optimal velocities to identify misleading velocity embeddings. Note that any conclusions are based on the underlying gene expression embedding, the significant vectors or the test-optimal velocities are the best representation based on this embedding but they can still have a low absolute test statistic (for example the Ngn3 low EP cluster, Fig. S3a,b).

For the dentate gyrus dataset [18] (Fig. 1d), the visualized velocities represent the high-dimensional data well, as can be seen from the non-uniform p-value distribution. The only exceptions are the isolated clusters where only a few vectors are significant, and the test-optimal velocities show some deviation. In addition, the connection of nIPC and radial glia-like cells is less faithfully represented, which could stem from the thin embedding structure in this region.

Wilk et al. [3] analyze the immune response to infections with SARS-CoV-2 (Fig. 1e) and generate a hypothesis based on the velocity embedding [3]. While we were not able to reproduce the same velocity embedding using the data and code provided by the authors, our overall layout of the expression data appears similar. The velocity visualization of this dataset has a poor representation of the original data: only a few velocity vectors are significant, the distribution of p-values is nearly uniform, and the test-optimal velocity differs vastly from scvelo’s embedding. Therefore, we suggest that the shown velocity embedding should not be used for the analysis of this dataset, and that further analysis should be performed based on the high-dimensional data.

We now turn to two alternatives for embedding RNA velocity datasets: Veloviz [14] and Nyström [15]. Veloviz first computes a gene expression embedding influenced by the RNA velocities and then uses scvelo’s algorithm to compute the velocity embedding. Nyström, on the other hand, uses the same gene expression embedding as scvelo, but computes a different velocity embedding. Comparing these methods on the pancreas data (with the stochastic velocity estimate), we get significant velocities for 5% of the data using veloviz, and the distribution of p-values is far from uniform (Fig. 2a). In contrast, the Nyström’s embeddings perform the worst of all three methods with only 3.2% significant vectors (Fig. 2b).

**Figure 2.**
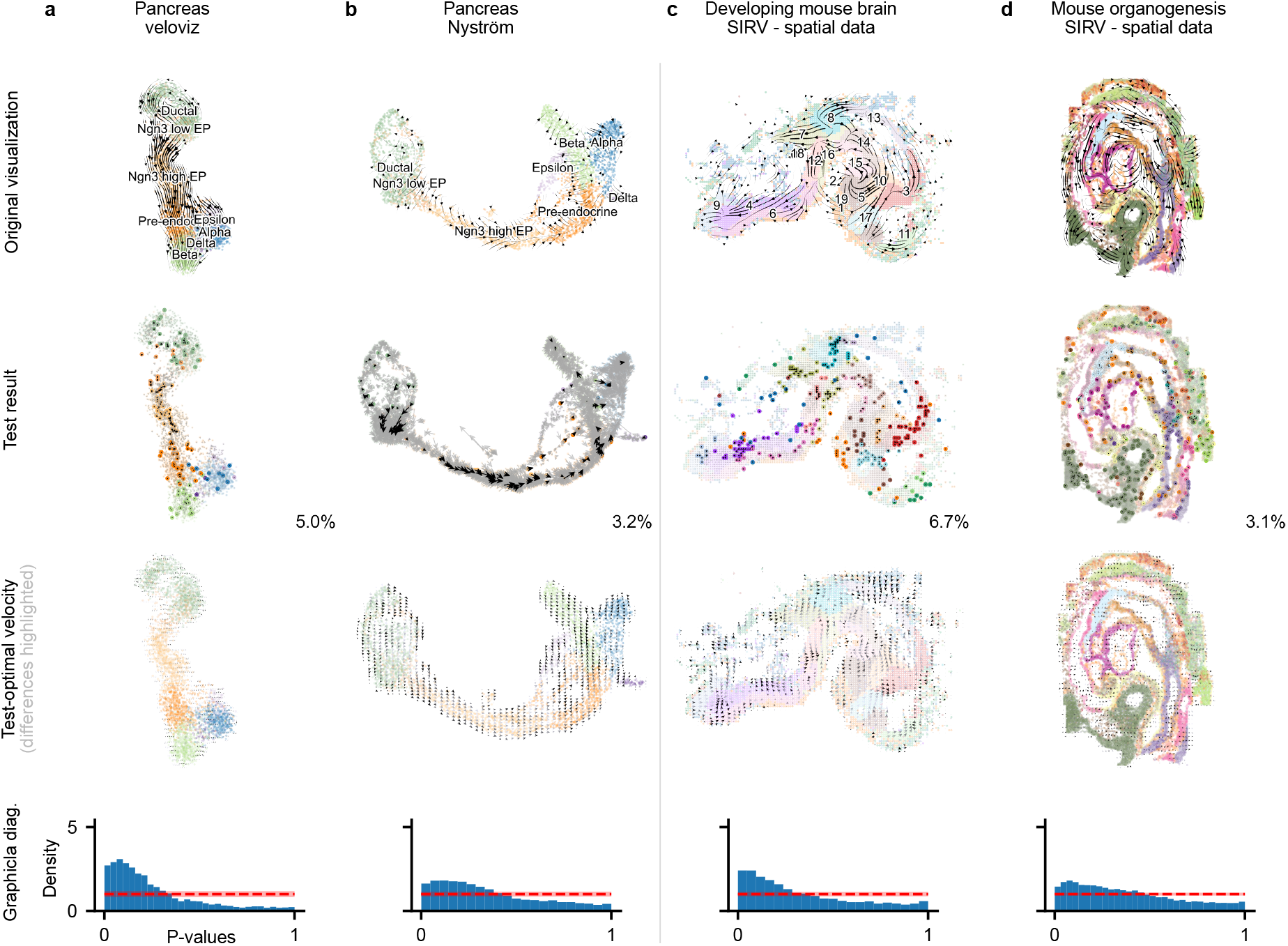
velotest for different velocity embedding methods and sequencing technologies. Panel rows organized as in Fig. 1b-e. velotest applied to **(a)** veloviz’ gene expression embedding of pancreas dataset combined with scvelo’s projection of velocity vectors, **(b)** Nyström embeddings on UMAP projection of pancreas dataset, **(c)** scRNA-seq data projected onto spatial transcriptomics measurements (HybISS) via SIRV of the developing mouse brain atlas, and **(d)** SeqFISH measurements of the mouse organogenesis dataset projected by SIRV.

We additionally apply velotest to spatial transcriptomic measurements. We use SIRV’s [22] projections of RNA velocities onto the spatial transcriptomics data of the developing mouse brain atlas [20] and mouse organogenesis dataset [21]. SIRV uses the x-y coordinates of the measurements as a replacement for the gene expression embedding of the scRNA-seq embedding methods and projects the simultaneously measured RNA velocity with scvelo’s algorithm onto these coordinates. For the developing mouse brain atlas, we observe a clustering of well and poorly represented velocities (Fig. 2c). This indicates that the cells in these well-represented clusters are developing into a state more similar to their physical neighbors. The mouse organogenesis dataset shows a weaker cluster structure (Fig. 2d), and the significant vectors are more uniformly distributed. Overall, the distribution of p-values is closer to the uniform distribution compared to most of scvelo’s embeddings, and calls for a more detailed evaluation of SIRV’s velocity embeddings.

## Discussion

We introduced velotest, a novel way of assessing the accuracy of RNA velocity visualizations by measuring how well the two-dimensional velocity embedding aligns with the high-dimensional velocity vector. Our experiments revealed large differences in embedding accuracies between different methods, datasets, and regions in a single dataset. velotest is neither another RNA velocity estimation procedure, nor does it assess the quality of the high-dimensional RNA velocity estimate (Fig. S1b). Instead, velotest enables the user to assess whether the hypotheses derived from inspection of velocity embeddings are consistent with the high-dimensional RNA velocity estimates. In summary, we envision velotest to enable reliable visualization-guided reconstruction of dynamic processes in future studies. It will allow users to identify misleading velocity embeddings and select those that accurately represent the high-dimensional RNA velocity. This will increase the likelihood of successful subsequent experimental validation of identified cell state transitions. The results using the test-optimal velocity also showed that there is a need for improved combined gene expression and velocity embedding methods as for some datasets even the best possible velocity did not represent the data very well.

We provide the Python package *velotest* ^1^ compatible with the widely used scverse’s anndata [23]. velotest can therefore directly be applied to any paired data with a corresponding embedding without further pre-processing and will provide an immediate quality check for embedded velocities.

## Methods

To apply *velotest*, we assume to have a high-dimensional dataset of gene expression levels and velocity vectors (**g, v**) ∈ (**G, V**) and corresponding two dimensional embeddings 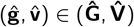. More specifically, 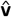 defines the vector pointing from **ĝ** to 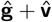.

### Testing procedure

To assess the quality of an embedded velocities 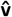 we quantify the velocity alignment using cosine similarity: First, we select neighbors of the cell in the direction of the velocity 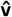 in the two-dimensional embedding space. Then, we evaluate whether the high-dimensional velocity **v** also points to those cells in the high-dimensional space using cosine similarity (Fig. 1a). The test statistic for 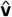 is then defined as the mean cosine similarity over all selected cells in the two-dimensional embedding space. We calculate this test statistic using all possible velocity vectors under *H*_0_ that are visually distinct from the embedded velocity (Fig. S2). To establish a notion of ‘visual distinction’, we introduce an exclusion angle *γ* which limits alternative velocities to be at least *γ* degree distinct from the embedded velocity 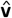. We conduct the test by calculating a p-value based on the test distribution under the null hypothesis for each cell individually, and correct via Bonferroni correction for multiple testing.

#### Test statistic

To compute the test statistic, we first identify the *k*-nearest neighbors NN(**ĝ**) in the two-dimensional space for every datapoint **ĝ** (default value *k* = 200, Fig. S4, see Choice of hyperparameters for a discussion). The test statistic *T* for a single data point 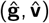 is then defined as

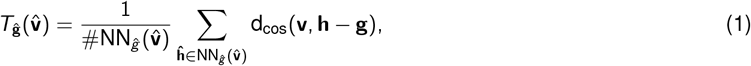

where 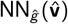 is a subset of NN(**ĝ**) which includes all gene expression embeddings which are within a 2*β* cone (with *β* being the neighborhood angle, Fig. 1a) of the visualized velocity 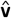. The cosine similarity d_cos_ is then calculated in the original, high-dimensional space. We take the preimage **v** of the visualized velocity 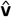 and the preimages **g** and **h** of the embedded gene expression levels and calculate 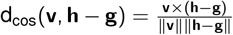. This statistic assesses whether the high-dimensional velocity **v** points in the direction of the preimage of the selected nearest neighbors 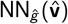.

As the distribution of this test statistic under the null hypothesis *H*_0_ does not have a closed form and will differ between cells, we infer the null distribution by evaluating the test statistic for alternative velocity embeddings 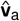 from the unit circle that are at least distinct from the visualized velocity by an angle of *γ* (see Fig. S5 for different choices of *γ*). To get 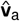, we first sample an angle *θ* ∈ [*γ*, 360°− *γ*) and convert it to its cartesian representation 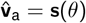.

As we have a finite neighborhood NN(**ĝ**), the test statistic *T*_**ĝ**_ can only take finitely many values and we can get the full mapping *T*_**ĝ**_: (0°, 360°) → (−1, 1), *θ* → *T*_**ĝ**_(**s**(*θ*)), in which we assume without loss of generality that 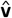 corresponds to an angle of 0°(Fig. S2).

With this mapping, we can calculate the p-value using the observed statistic 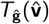 as

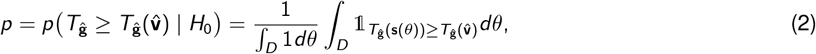

where the integration domain *D* ⊆ [*γ*, 360°− *γ*] includes all valid alternative angles that are visually distinct from the embedded velocity 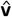 (see Alg. 1 for the complete testing procedure).

We perform this test for every cell in the dataset, typically in the order of thousands for scRNA-seq datasets. We therefore correct for multiple testing to avoid Type I errors (false positives). As the tests have a non-trivial dependence structure, we resort to correcting via the Bonferroni correction [17] despite its conservatism, as it does not have any dependency assumptions (unlike the Benjamini-Hochberg procedure [24]). The Benjamini–Yekutieli procedure [25] (which also relaxes the assumptions on dependency between tests) yields qualitatively similar results to the Bonferroni correction. Having said that, even for non-corrected test results, the overall interpretation vastly remains the same (Fig. S6), e.g., for the Pancreas dataset, the alpha and epsilon cluster are still not significant.

##### Algorithm 1

Test for single cell (*g, v*) if its embedded velocity 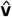 is better than a randomly chosen velocity.

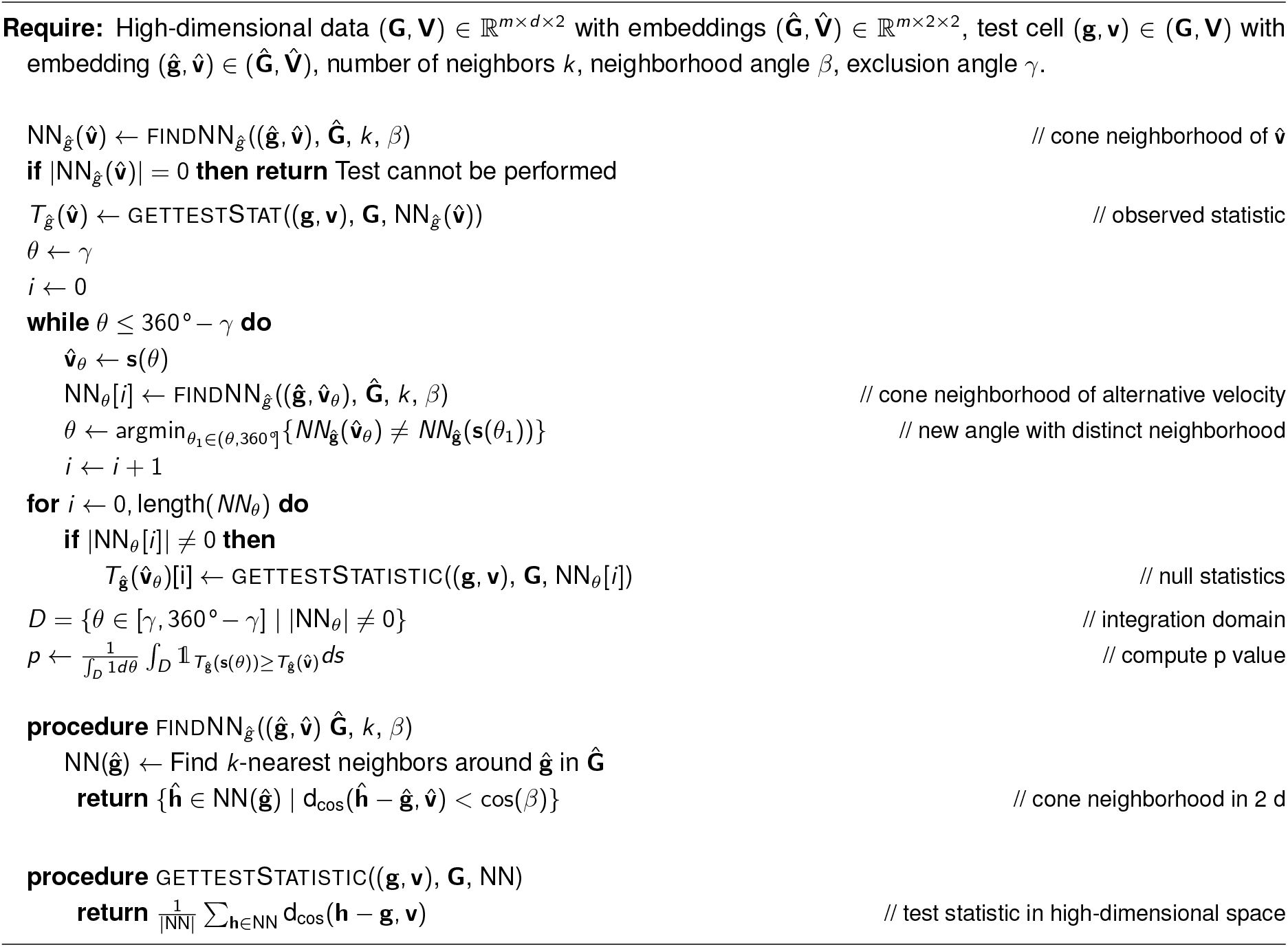

#### Isolated velocities

As our test statistic is based on neighboring cells that lie within a 2*β* cone, we cannot compute a test statistic for cells 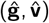 for which the cone neighborhood 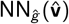 is empty. We therefore exclude these cells from our test procedure, which typically leads to very few untested cells on the boundary of the visualized embeddings (Fig. S7). Similarly, we exclude velocities from *D* that have empty neighborhoods from the computation of the p-value, and the integration domain *D* becomes finally *D* = {*θ* ∈ [*γ*, 360°− *γ*] | NN_*ĝ*_(**s**(*θ*)) *≠* ∅}. The latter is especially prominent for embeddings that have thin structures, like the *developing neutrophil* cluster in the Covid dataset (Fig. 1e). For such structures, the support of the null distribution is strongly reduced as alternative velocities need non-empty cone neighborhoods.

#### Connection between significance level *α* and exclusion angle *γ*

Using the exclusion angle *γ* to limit the evaluation domain of the test statistic 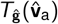 has an effect on the significance level *α* for our testing procedure (regardless of the correction for multiple testing). Considering a significance level *α, α ×* 100% of the overall velocities are allowed to have a higher test statistic than the observed statistic. On the other hand, for an exclusion angle *γ* we additionally allow alternative velocities in a cone of 2*γ*, which are up to 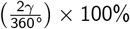 of the overall velocities, to have a higher test statistic than the observed statistic. With both significance level *α* and exclusion angle *γ* in play, we have up to a combined 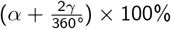 effective samples of the overall velocities. Obtaining this amount of effective samples is equivalent to considering an effective significance level of up to 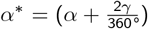 for the decision criteria on our testing procedure. Thus, the empirical relation between a chosen significance level *α* and exclusion angle *γ* effectively imposes an updated significance level *α*^∗^ for our tests. However, if all alternative velocities do not produce a higher test statistic, the effective significance level is not increased as much, and under the specific case where none of the test statistics for the alternative velocities in the exclusion cone are higher than the observed statistic, the effective significance level stays at *α*^∗^ = *α*.

#### Judging the overall embedding instead of individual velocity vectors

Multiple testing procedures require correction methods to adjust significance levels for sequential tests. This directly affects the decision criteria of rejecting the null hypothesis based on the resulting p-values. While multiple testing is necessary to answer questions regarding the statistical significance of observed velocities at the single-cell level, we provide an alternative graphical diagnostic to tackle inference for the observed velocities for the overall data.

We consider the p-values computed based on the test statistic 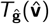 such that *p*_*ĝ*_ is the p-value for the cell associated with **ĝ**. We further consider *p*_*ĝ*_ to be realizations of the random variable *P* associated with the p-values. The random variable *P* is therefore uniformly distributed under the null hypothesis, *P* ∼ *U* (0, 1) (see Murdoch et al. [26] for more details). This follows from the p-values being a function of the test statistic *T* such that *P* = *F*_*T*_ (*T*). Then, the cumulative distribution function (CDF) of *P* can be derived using the probability integral transform of the test statistic *T* under the null such that *F*_*P*_ (*p*) = *p*. This is the CDF of a continuous uniform distribution over the interval [0, 1].

Thus, by plotting the histogram of samples *p*_*ĝ*_, we can graphically identify deviations from uniformity as evidence in favor of rejecting the null hypothesis. For the visualization, we allocate the p-value samples to histogram bins. This allocation follows a binomial distribution based on the number of p-value samples and the proportion of allocating a p-value to a bin. We use Freedman-Diaconis rule [27] to choose the number of bins and construct a 95% confidence band based on the resulting binomial distribution. Any deviations outside of this confidence region are directly attributed to evidence against the null distribution. For a specific dataset, we interpret the non-uniformity as evidence against the visualized velocity being a random velocity embedding. Based on our experiments, the graphical diagnostic suggests strong evidence against the visualized velocity being just a random velocity embedding (through departure from uniformity) for the pancreatic endocrinogenesis dataset using the dynamic model (see Fig. 1c). On the other hand, we lack sufficient evidence to make the same statement with a similar level of confidence regarding the velocity embedding of the SARS-CoV-2 data [3] (Fig. 1e).

### Test-optimal velocities

Using the above-defined test statistic 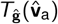 for alternative velocities, we can search for velocities that preserve the information of the high-dimensional velocity **v** as much as possible. To this end, we solve

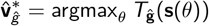

to find the velocity embedding which would maximize the test statistic given gene expression embeddings **Ĝ**. While this alternative velocity embedding is optimal under the given gene expression embedding and test statistic, we want to emphasize that the preserved information of the high-dimensional velocity might still be limited. This is indicated by a test statistic that is close to 0 (or negative), representing that on average the high-dimensional velocity **v** is perpendicular (or pointing into an opposite direction) to the preimages of the cells in the 2*β* cone and their corresponding high-dimensional vector **h** − **g** (Fig. S3). This can either stem from the underlying gene expression embedding or the data itself.

### Choice of hyperparameters

The results of velotest rely essentially on three hyperparameters: the neighborhood size *k*, the neighborhood angle *β*, and the exclusion angle *γ*. Their influence and optimal choice are dependent on the data as well as the embedding structure.

We recommend choosing the neighborhood size *k* as big as possible without spanning unconnected areas on the embedding (see Fig. S4b,c for *k* = 200). By spanning unconnected regions, the test statistics for alternative embedding vectors can be unreliable as commonly used embedding procedures for scRNA-seq (e.g., t-SNE) preserve local structures but less of the global arrangement [28]. For the presented datasets, we chose a default value of *k* = 200, reduced it to 100 for the dentate gyrus and mouse organogenesis datasets (Fig. S4b,c), and increased it to 300 for the gastrulation erythroid dataset after visual inspection.

The two angular parameters *β* and *γ* are dependent on the visual interpretation: the neighborhood angle *β* defines which gene embeddings are accepted to be in a similar direction as the visualized velocity 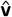. We chose *β* = 22.5°as a default value, and therefore interpret all gene embeddings within the 2*β* = 45°cone to be aligned with 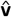. Choosing too small values would lead to a test statistic that is very sensitive to the exact placement of the cell embeddings. On the contrary, choosing too large values would violate the notion that those neighbors lie in the direction of the velocity vector. We found robust results within a reasonable range of *β* = 5°to *β* = 30°(Fig. S8).

The exclusion angle *γ* has a high influence on the test results. For *γ* = 0°, we get the strictest test, which often leads to no significant velocities after correction for multiple testing (Fig. S5, first row). With an increasing exclusion angle *γ*, our test gets less strict (see discussion above). If some or all of the test statistics for the velocities in the exclusion angle are larger than the observed test statistic (Fig. S2a), the p-value would increase if the exclusion cone were not to be used. But those alternative velocities are, in fact, visually hard to distinguish from the embedded velocity. However, for a lot of cells, the statistics in the exclusion angle lie on both sides of the observed test statistic, and these cells would become significant only for very large exclusion angles (Fig. S2b,d and Fig. S5). Despite the influence of this hyperparameter, we see similar patterns for a broad range of exclusion angles *γ* (Fig. S5). For example, even for very large exclusion angles, the alpha and epsilon clusters in the pancreas dataset do not show good embedding qualities for the stochastic velocity model, similarly for the bone marrow dataset.

Additionally, while we recommend choosing *γ* as small as possible, we have to bear in mind what we can actually differentiate as visually distinct, and more importantly, how we further interpret the visualization. Especially for stream plots (e.g. Fig. 1b, first row), but even for grid plots (e.g. Fig. 1b, last row), multiple velocity vectors are summarized, and one is often interested in the overall direction of the trajectory rather than the exact angle of one specific cell. We name the test velotest_*γ*_ based on the used exclusion angle *γ* to make clear how strict the applied test was, e.g., velotest_10_ for *γ* = 10°.

### Runtime of velotest

The runtime of velotest is mainly dependent on the number of datapoints in the datasest. For the investigated datasets we find reasonable runtimes of velotest_10_ below one minute up to two minutes on a M1 Pro MacBook and 32GB memory based on 300 neighbors and *β* = 22.5 (Tab. 1).

**Table 1.** Dataset sizes and runtimes of velotest.

| Dataset | # Cells | # Velocity Genes | Total runtime [min] |
| --- | --- | --- | --- |
| Pancreas | 3696 | 1038 | 0:33 |
| Covid | 3076 | 126 | 0:25 |
| Dentate gyrus | 2930 | 557 | 0:24 |
| Bone marrow | 5780 | 452 | 0:45 |
| Gastrulation erythroid | 9815 | 487 | 1:26 |

## Acknowledgements

This work was funded by the Deutsche Forschungsgemeinschaft (DFG, German Research Foundation) under Germany’s Excellence Strategy (EXC number 2064/1, Project number 390727645), Collaborative Research Center 391 (Spatio-Temporal Statistics for the Transition of Energy and Transport, 520388526), and by the German Federal Ministry of Education and Research (BMBF): Tübingen AI Center, FKZ: 01IS18039. The authors thank the International Max Planck Research School for Intelligent Systems for supporting S.B. and S.M. The authors thank Julius Vetter, Jan Schleicher, and Leonie Kolmar for feedback on an earlier version of this manuscript.

## Author Contribution

Conceptualization: S.B., C.S., Formal analysis: S.B., C.S., S.M., M.C., Investigation: S.B., P.G.P. Methodology: S.B., C.S., Software: S.B., Supervision: C.S., M.C., J.H.M., Visualization: S.B., C.S., Writing – original draft: S.B., C.S., Writing – review & editing: all authors, Funding acquisition: M.C., J.H.M.

## Supplementary Text and Figures

### Additional Datasets

Besides the previously presented datasets, we also analyzed the bone marrow dataset [19] and gastrulation erythroid dataset [18] with velotest.

The visualized velocities of the bone marrow dataset [19] are overall poorly representing the high-dimensional data (Fig. S1a), as indicated by only few significant cells (2.2%) and a relatively flat p-value distribution. Additionally, the test-optimal velocity embeddings are not aligned with the visualized ones, and actually point away from the embedded expression data. Similar to Bergen et al. [12], we also find problematic visualizations in the erythroid clusters (Ery 1, Ery 2). For these clusters, Gao et al. [29] have shown that using different velocity estimates (UniTVelo [29]) results in a change of direction matching the expected trajectory.

The gastrulation erythroid dataset [18] (Fig. S1b) exhibits small gaps in the distribution of significant velocities, which coincide with patterns in the test-optimal velocity. It has a non-uniform p-value distribution, but it is less distinct than for the pancreas or dentate gyrus dataset. Note that this finding is again independent of the problematic high-dimensional velocity estimates, as pointed out for this dataset by Bergen et al. [12].

**Figure S1.**
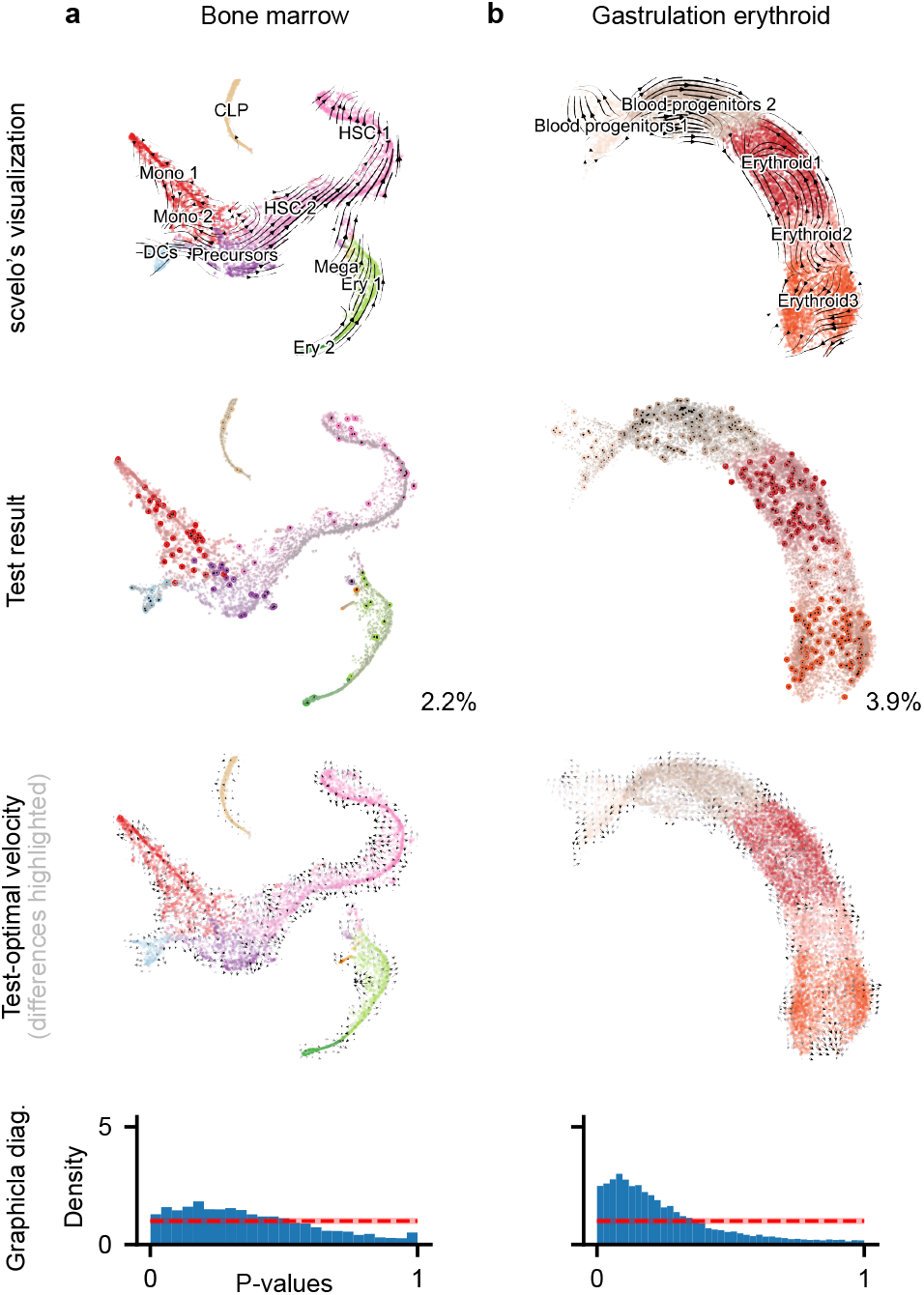
velotest for additional datasets. Panel rows organized as in Fig. 1b-e. velotest applied to scvelo embeddings of **(a)** the bone marrow dataset [19], and **(b)** the gastrulation erythroid dataset [18]. The gastrulation erythroid dataset is a good illustration that velotest only tests the faithfulness of the embedding and cannot be used to judge the high-dimensional estimates, as the distribution of p-values is far from uniform even though the velocity is estimated wrongly [12]

**Figure S2.**
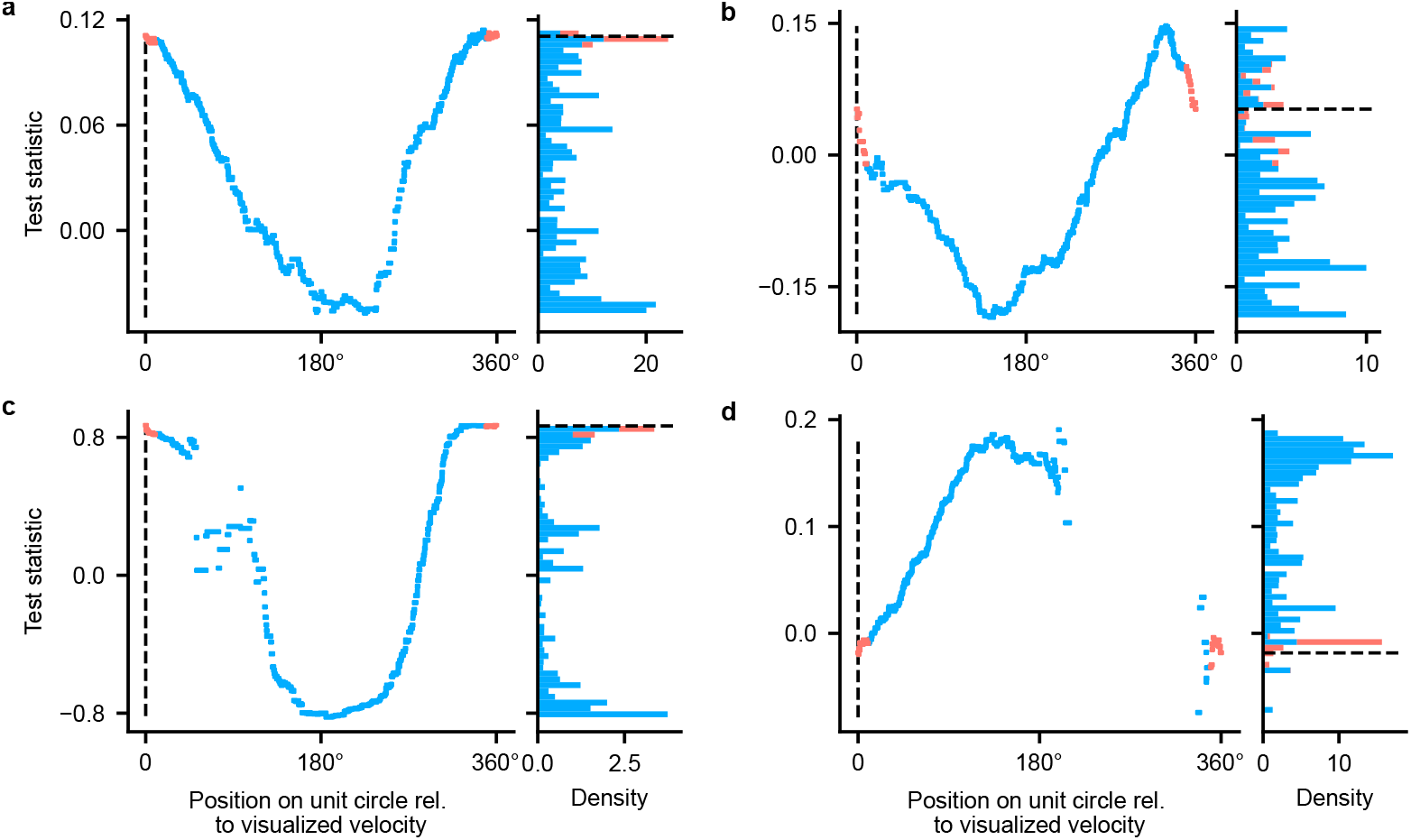
Example test statistic *T*_ĝ_. *Left plots:* Test statistic for velocity embedding and alternative velocity vectors across the full unit circle. Positions inside the exclusion cone *±γ* are marked in red. *Right plots:* Density over all possible test statistics for the chosen cell. Test statistics for positions inside the exclusion cone are marked in red. The test statistic of the visualized velocity is marked by a black dashed line. **(a)** Cell with a large test statistic near the maximum 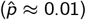. **(b)** Cell with a small test statistic, which is not significantly different from random directions 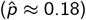. **(c)** Cell where the visualised velocity has the highest test statistic, which makes it significant 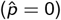. **(d)** Cell where test statistic is not defined for full domain [0°, 360°] 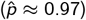.

**Figure S3.**
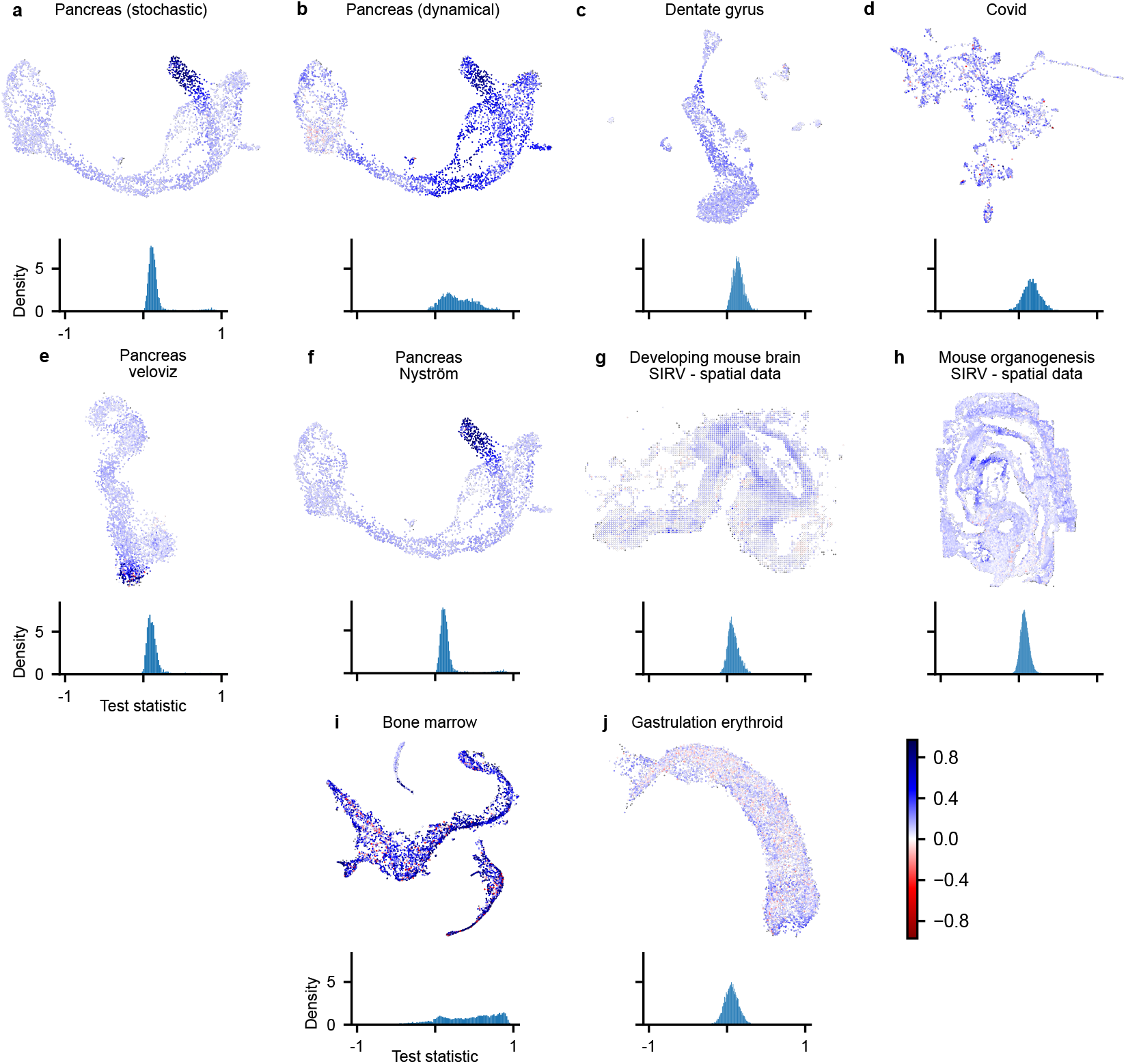
Test statistics for test-optimal velocities. **(First, third, and fifth row)** Test statistic for the optimized velocity embedding 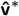 across all datasets. **(Second, fourth and sixth row)** Histogram of test statistics in corresponding scatter plot. **(a**,**b)** The UMAP of the gene expression data generally seems to be able to better represent velocities from the dynamical model than the stochastic model. The pre-endocrine and endocrine (alpha, beta, delta, and epsilon) clusters are well represented for the dynamical velocity, whereas only the beta cluster is well represented for the stochastic velocity. **(b**,**i)** The pancreas dataset with velocities from the dynamical model as well as the bone marrow dataset have a very broad distribution of test statistics. **(b**,**d**,**g-j)** Even the highest possible test statistic given this embedding, produced by the test-optimal velocities, is negative for some cells. A cosine similarity of zero indicates that the high-dimensional velocity vector of the tested cell and the vector towards the neighbor (to which the velocity embedding points) are perpendicular. A negative cosine similarity means that it points away from the neighbor. **(a**,**e**,**f**,**h)** The visualizations of the pancreas dataset with velocities from the stochastic model as well as the mouse organogenesis dataset have clusters of test-optimal velocities with a high test statistic.

**Figure S4.**
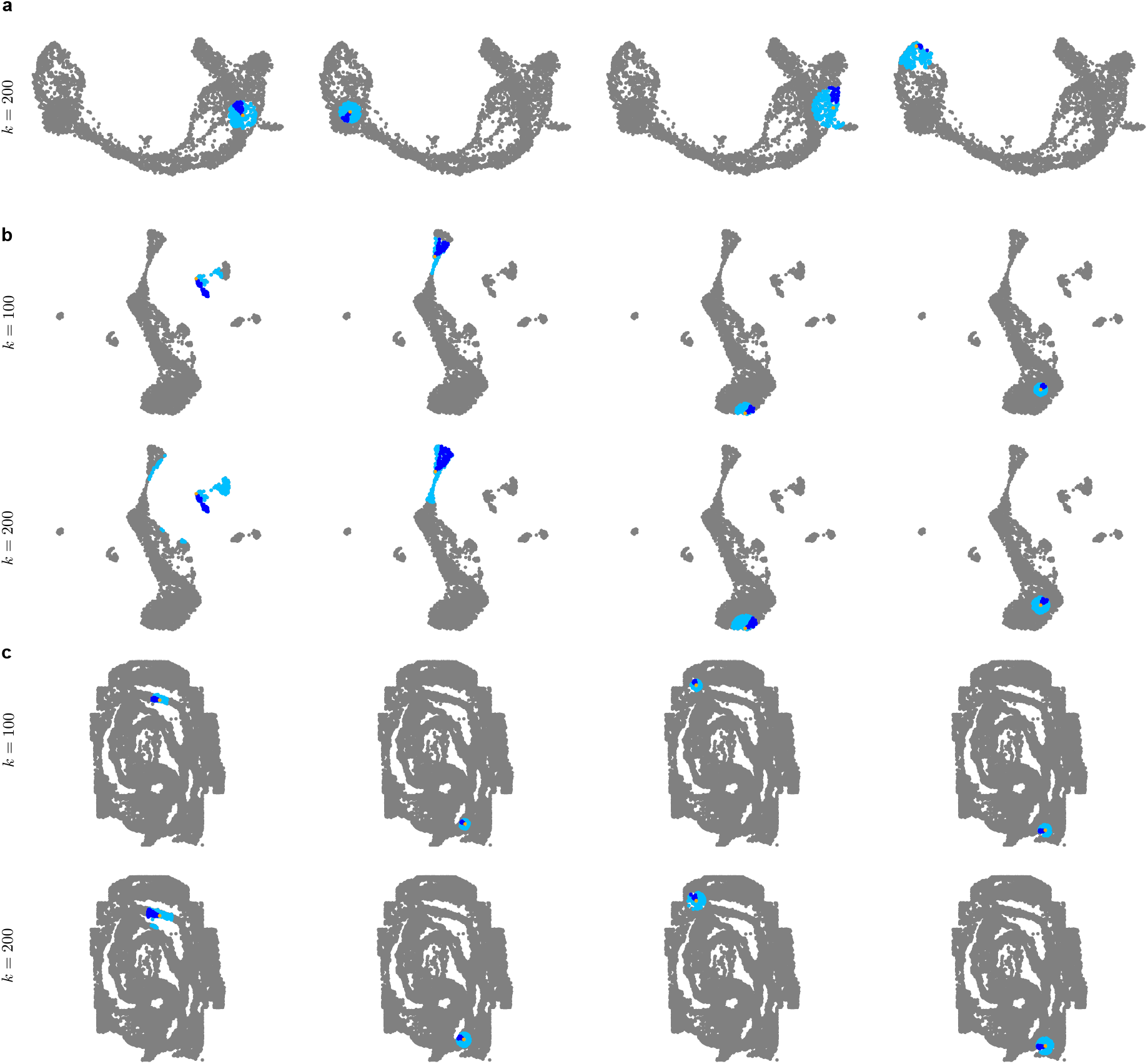
Example neighborhoods with varying *k*. Full neighborhoods NN_ĝ_ (light blue) and cone neighborhoods 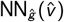 (dark blue) for the visualized velocity 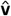 for four example gene embeddings **ĝ** (orange) from **(a)** the pancreas dataset, **(b)** the dentate gyrus dataset, and **(c)** the mouse organogenesis dataset.

**Figure S5.**
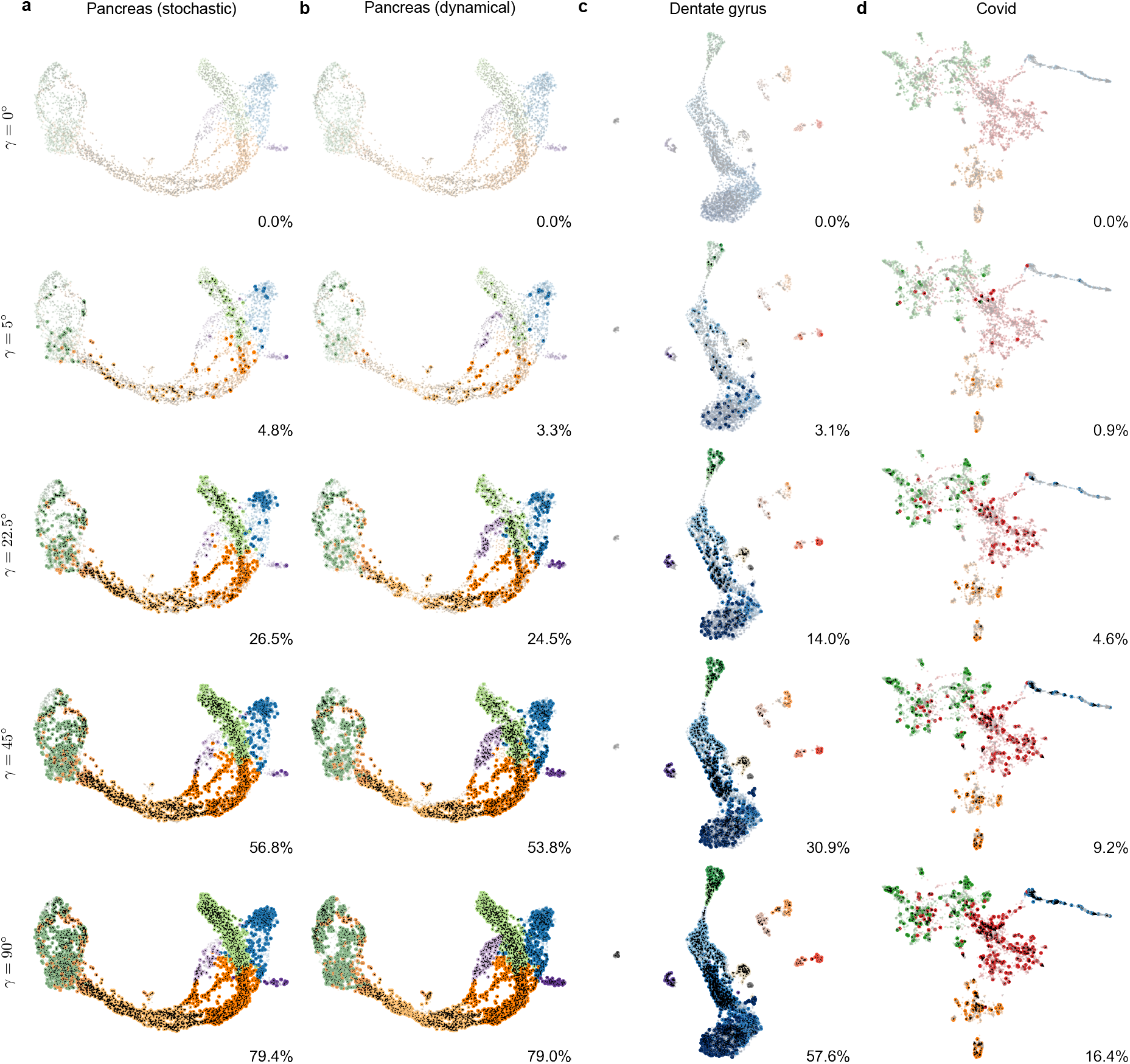
velotest_*γ*_ with varying exclusion angle *γ*. Applying velotest with no exclusion angle (0°), and increasing angles of 5°, 22.5°, 45°and 90°.

**Figure S6.**
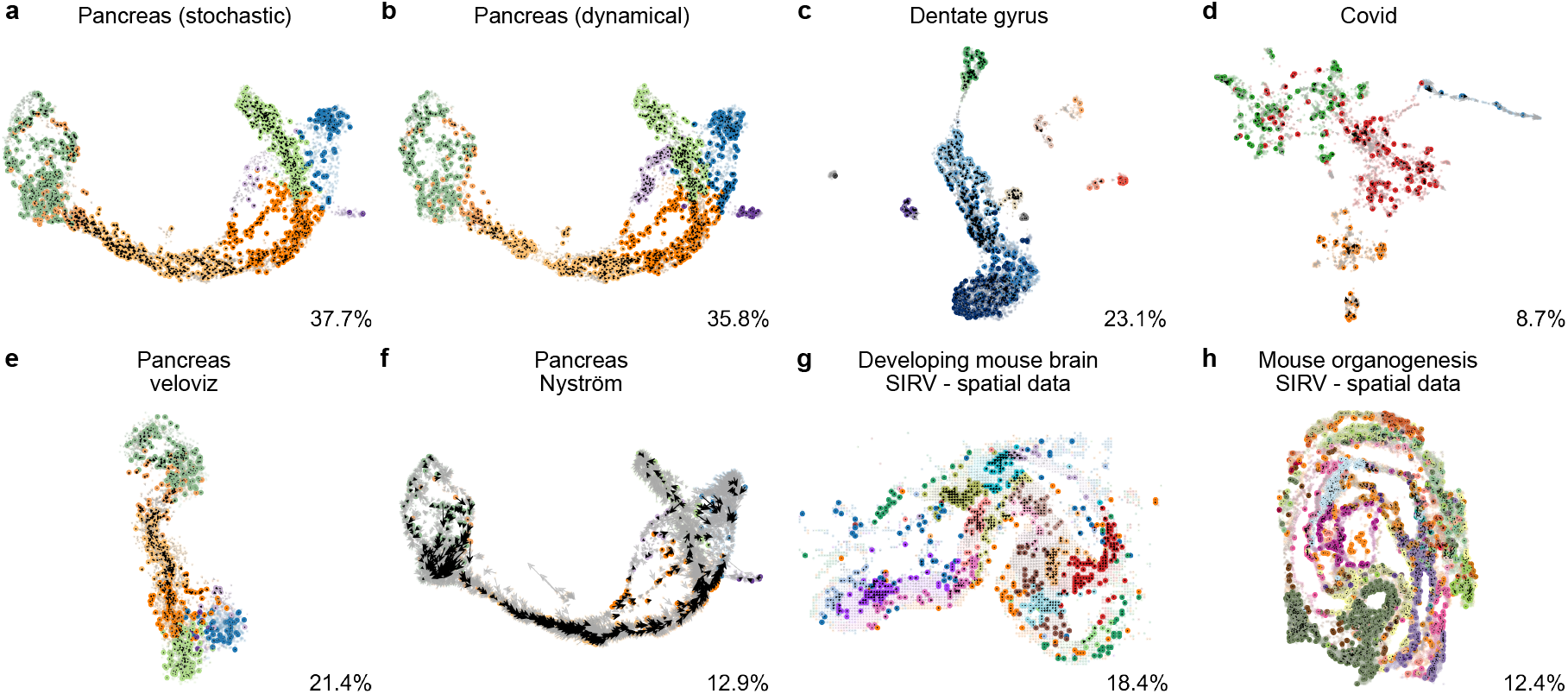
velotest_10_ without correcting for multiple testing. The null hypothesis is rejected for significantly more tests compared to Fig. 1 (with Bonferroni correction).

**Figure S7.**
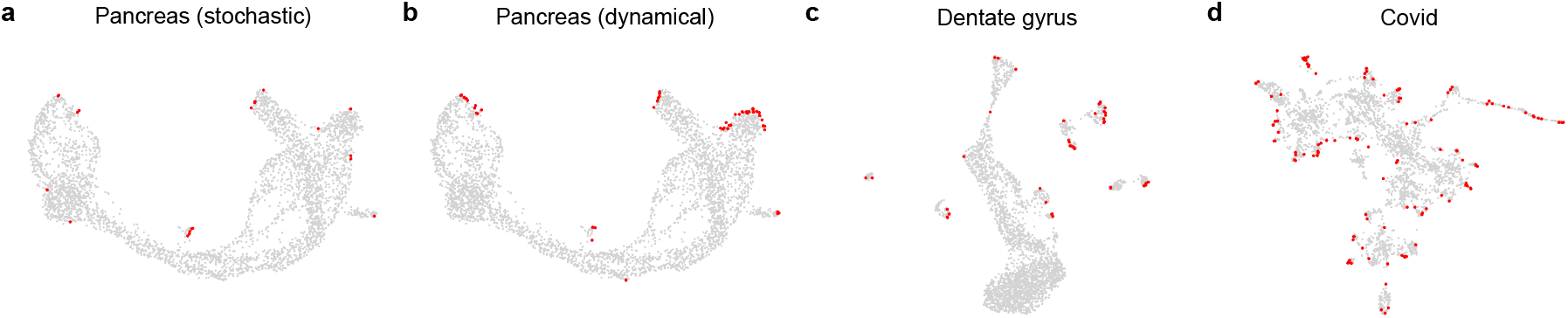
Untested velocities. Cells for which the neighborhood of the visualized velocity 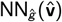 is empty cannot be tested. These cells (marked in red) typically lie on the boundary of the visualization. We get *n* = 20 (≈ 0.54%), *n* = 51 (≈ 1.38%), *n* = 42 (≈ 1.43%), and *n* = 119 (≈ 3.87%) of untested velocities for the embeddings a) to d).

**Figure S8.**
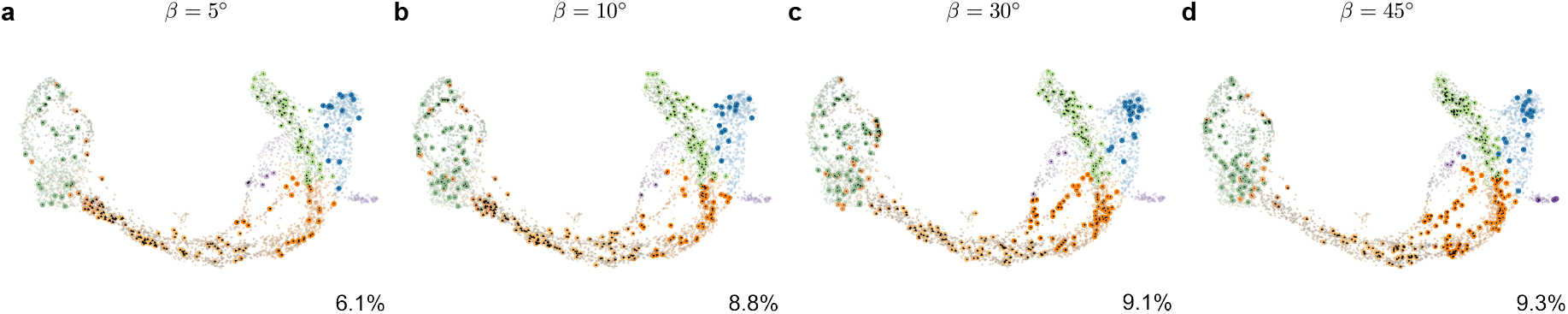
velotest_10_ with varying cone angles *β* for the Pancreas dataset. The results are consistent between *β* = 5°and *β* = 30°, with fewer significant vectors for *β* = 5°. However, for a large angle of *β* = 45°, there are local changes, e.g., the beginning of the *Ngn3 high EP* cluster (left part of yellow cluster) contains barely any significant vectors.

## Footnotes

1 https://github.com/mackelab/velocity-hypothesis-test/

